# Optimized LC-MS/MS method for Doxorubicin quantification: validating drug uptake and tumor reduction in zebrafish xenograft model of breast cancer

**DOI:** 10.1101/2024.08.09.607268

**Authors:** Ghazala Rahman, Atanu Pramanik, Susmita Das, Anindya Roy, Anamika Bhargava

## Abstract

Doxorubicin, a potent chemotherapeutic drug, is widely used against various cancers, notably breast cancer. While its efficacy is well-documented, precise dosage determination in experimental models remains challenging. Zebrafish xenografts of various cancers confirm doxorubicin’s anti-cancerous effect; however, since doxorubicin treatment of zebrafish larva is done by adding doxorubicin to fish water, the precise chemotherapeutic dosage for zebrafish larva remains unknown. In this study, we provide a liquid chromatography tandem mass-spectrometry (LC-MS/MS) method for quantifying doxorubicin uptake in zebrafish larvae and thus provide a direct estimate of doses required for the therapeutic effect. Alongside quantification, we measured the therapeutic effect of doxorubicin in zebrafish larvae xenografted with triple negative breast cancer cell line, MDA-MB-231. LD_50_ value of doxorubicin was first determined by incubating 3-days post fertilization (dpf) larvae with different doses of doxorubicin for 72 h. Doxorubicin was quantified both from zebrafish larval homogenate and treatment solution. Analysis was performed by selected-reaction monitoring (SRM) scans in positive ionization mode. LD_50_ value for 72 h calculated to be 35.95 mg/L. As expected, doxorubicin-treated xenografts exhibited a significant reduction in tumor growth. The range of limit of detection (LOD) and limit of quantification (LOQ) for doxorubicin were 2 and 5 μg/L respectively. Intra- and inter-day accuracy was within the range of 82-114%. Overall, in this study we describe a reliable method for quantifying doxorubicin in zebrafish larvae. Our study facilitates precise dosage estimation, enhancing the relevance of zebrafish xenograft model in cancer research and potentially improving translational applications of chemotherapeutic treatments.

## Introduction

Chemotherapy still remains a cornerstone in modern oncology. Chemotherapy employs various chemotherapeutic drugs that broadly function by targeting and disrupting crucial processes essential for cell survival and cancer progression (1). Globally, the leading cancer incidence is from breast cancer that is being largely targeted by doxorubicin, a widely known chemotherapeutic drug belonging to anthracyclines group (2). In addition, doxorubicin, is also used against other types of cancers as well including bladder cancer (3), lung cancer (4) and leukemia (5). It functions by inhibiting topoisomerase II thereby disrupting DNA replication and transcription (6).

Previous studies have shown the anti-cancerous efficacy of doxorubicin in both *in-vitro* (7,8) and *in-vivo* models (9,10). Among the *in-vivo* models, zebrafish xenografts have been extensively used for high throughput screening of potential anti-cancerous compounds (11–13). Notably, breast cancer zebrafish xenograft model also helps us to understand cancer physiology and to discover novel targets (14,15). Drug exposure to zebrafish xenografts are simply through fish water. Bio-uptake quantification of doxorubicin has been carried out by Igartua et al.,2018 through fluorescence imaging by virtue of its intrinsic fluorescence property (16). However, this doesn’t provide absolute quantification. A more reliable approach involves mass spectrometry, which offers accurate quantification of compound uptake within the larval body. Although the LC-MS/MS method has been developed and validated for numerous compounds (17,18), the quantification parameters for doxorubicin uptake in zebrafish larvae are yet to be determined and is the subject of this study.

In this study, we describe a LC-MS/MS method to determine the precise doxorubicin uptake from the fish water and in the zebrafish larval homogenate, facilitating precise dosage estimation.

## Materials and Methods

### Animal cell culture

Human breast cancer cell line, MDA-MB-231 was procured from National Centre for Cell Sciences (NCCS, Pune, India) and maintained in Dulbecco’s Modified Essential Medium (DMEM, P4333, Sigma-Aldrich) supplemented with 10% fetal bovine serum (RM9955-500 mL, Himedia), 50 U/mL Penicillin-streptomycin (15140-122, Gibco) at 37 ⁰C in 5% CO_2_ in a humidified chamber. MDA-MB-231 cells were labelled with Vybrant Dil cell labelling solution (V22885, Invitrogen) according to manufacturer’s guidelines for microinjection.

### Zebrafish housing and maintenance

Adult wild type zebrafish were purchased from local commercial suppliers and housed in a standard 14:10 h light and dark photoperiod with temperature maintained at 28 ⁰C. Fishes were fed twice a day. Adult fishes (female: male-2:1) were kept overnight for mating in a breeding chamber and an average of 100-300 larvae were obtained per chamber. Fertilized healthy larvae were screened at 5-hour post fertilization (hpf) and maintained in 1 X E3 media with 0.2 mM N-Phenyl Thiourea (PTU) (P7629-10G, Sigma-Aldrich) at 28 ⁰C in a BOD incubator. PTU was used to inhibit pigmentation for maintaining larval transparency for imaging.

### Determination of doxorubicin LD_50_ in zebrafish larvae

In this study, MDA-MB-231 xenografted zebrafish were treated with doxorubicin after 24 hours of injection at 3-days post fertilization (dpf) for 3 days, therefore, zebrafish larvae were treated with the same regime for determining the doxorubicin LD_50_ value. For this, zebrafish larvae at 3 dpf were randomly selected and 5 larvae per well were manually transferred to a flat 24-well plate. Each treatment was performed in three technical replicates. Doxorubicin hydrochloride (D4193, Tokyo chemical industry, TCI) of 0.1, 1, 10, 25, 50, 100 mg/L concentrations were prepared from a stock of 5000 mg/L in 1 X E3 media supplemented with 0.2 mM PTU. Larvae from each well were incubated in 400 μL of respective treatment solutions for three days in dark conditions to avoid photodegradation of doxorubicin. Similar to zebrafish xenografts, these larvae were incubated at 34 ⁰C which was optimised for tumor growth in xenografts. Mortality was assessed and dead larvae were removed every 24 h. Morphological abnormalities such as yolk sac edema, pericardial edema, tail bending and bent spine were recorded at 3-days post treatment (dpt). Percent mortality and abnormalities were calculated and LD_50_ value was determined for 3 dpt using the equation Y=100/(1+10^((LogIC50-X)*HillSlope)) by GraphPad Prism software version 8.0.2. Three biological replicates were performed for each concentration.

### LC-MS/MS method development for quantification of doxorubicin uptake by zebrafish larvae

#### Doxorubicin exposure to zebrafish larvae and sample preparation

Thirty zebrafish larvae at 3 dpf were treated with 3 ml of doxorubicin solution (5 mg/L and 10 mg/L) prepared in 1 X E3 media with 0.2 mM PTU. These concentrations were chosen as no significant mortality was observed in the doxorubicin larval toxicity study for these concentration groups. Control larvae were incubated only with 1 X E3 media containing 0.2 mM PTU. All the zebrafish larvae were incubated in the BOD incubator at 34 ⁰C in dark condition for 3 days. After 3 dpt, each group consisting of 30 larvae were first centrifuged to separate the supernatant (only incubation media) and pellet (only larvae). Larvae pellet was then washed thrice with deionised water and homogenized for 1 min using hand held homogenizer in 500 μL of deionised water. Homogenate was centrifuged at 10,000 rpm for 10 mins. Supernatant was filtered with 0.22 μm syringe filter. The concentration of internal standard, colchicine was maintained at 250 μg/L. Internal standard is a stable, non-interfering compound with similar retention time with the analyte which helps in accurate and precise quantification of the analyte. Drug uptake per larva was calculated using the following formula [ng/larva = concentration (ng/mL) X 0.5 mL /No. of larvae in the sample] as mentioned by Zhang et al., 2015 (18).

#### Reagents and solution preparation for LC-MS/MS

The reference standard, doxorubicin hydrochloride (purity >95%) was obtained from Tokyo chemical industry (TCI, D4193). The internal standard colchicine was purchased from Sigma (C9754). ULC-MS grade acetonitrile (0001204104BS) and methanol (0013684104BS) were supplied by Biosolve, formic acid (98+%) by Thermoscientific (147930010) and deionised water by Millipore (7732-18-5). Stock solutions of 1 mg/mL doxorubicin and colchicine were prepared by dissolving 10 mg of each in 10 mL Water: Methanol (50:50) separately. Each stock solution was stored at -80 ⁰C until further use.

#### Preparation of calibration samples for LC-MS/MS

Nine hundred zebrafish larvae of 6 dpf were washed twice with deionised water and homogenised in 15000 μL of deionised water to be used as matrix in calibration curve preparation. Doxorubicin stock solution of 1 mg/mL (10,00,000 μg/L) was diluted to a primary concentration of 0.05 mg/mL (50,000 μg/L) which was serially diluted to 20000, 10000, 5000, 2000, 1000, 500, 200, 100, 50, 20, 10 μg/L. The standard solutions were further diluted 10 times by mixing 50 µL standard solution to 50 µL of internal standard (2500 µg/L colchicine) with 200 µL of matrix and 200 µL solvent [Water: Methanol (50:50)] and analyzed on LC-MS/MS system. The resultant concentrations of doxorubicin were 1, 2, 5, 10, 20, 50, 100, 200, 500, 1000, 2000, 5000 µg/L and the peak area values obtained from the LC-MS/MS analysis were used for plotting the linear calibration curve.

#### Instrumentation for LC-MS/MS

The LC system was a Thermo Ultimate 3000 with autosampler (Thermo scientific). The mass spectrometer used was a Thermo TSQ Altis (Triple Quadrupole) (Thermo scientific) equipped with Heated Electrospray Ionization (HESI), operated in a positive ion mode, and analysis was carried out in Selected reaction monitoring (SRM) scan modes. The autosampler, tray and column temperatures were maintained at 7 ⁰C, 10 ⁰C and 35 ⁰C respectively. Table 1 shows the chromatographic conditions. MS conditions including global and scan parameters are listed in Table 2 and 3 respectively.

**Table 1:** HPLC conditions.

|  |  |
| --- | --- |
| Solvent A of mobile phase | Water with 0.1% Formic Acid |
| Solvent B of mobile phase | Acetonitrile: Methanol (800:200) with 0.1% formic acid |
| Flow (µL/min) | 300 |
| Column | Hypersil Gold column (100x2.1x1.9µm) |
| Mobile-phase gradient | 0-15 min 75% B, 15.1-16 min 75% B, 16.1-18 min 10% B |

**Table 2:** MS condition: Global Parameters.

| Global Parameters |  |
| --- | --- |
| Spray Voltage | Static |
| Positive Ion (V) | 3500 |
| Sheath Gas (Arb) | 50 |
| Aux Gas (Arb) | 10 |
| Sweep Gas (Arb) | 0 |
| Ion Transfer Tube Temperature (°C) | 325 |
| Vaporizer Temp (°C) | 350 |

**Table 3:** MS condition: Scan parameters.

| SRM Properties |  | SRM Table |  |
| --- | --- | --- | --- |
| Cycle Time (sec) | 0.8 | Doxorubicin Precursor ion<br>(m/z) | 544.17 |
| Q1 Resolution (FWHM) | 0.7 | Doxorubicin Product ion (m/z) | 361.12,<br>397.10 |
| Q3 Resolution (FWHM) | 1.2 | Doxorubicin Collision Energy<br>(V) | 18, 20.12 |
| CID Gas (mTorr) | 1.5 | Colchicine Precursor ion (m/z) | 400.08 |
| Source fragmentation (V) | 0 | Colchicine Product ion (m/z) | 310.16,<br>358.16 |
| Chromatographic Peak Width<br>(sec) | 6 | Colchicine Collision Energy<br>(V) | 25.77,<br>21.18 |

#### LC-MS/MS Method validation for doxorubicin quantification

Twelve different standard solutions spiked into control (blank) zebrafish larval homogenate were used to construct the calibration curve. The linear ranges of doxorubicin taken were 1-5,000 μg/L (1, 2, 5, 10, 20, 50, 100, 200, 500, 1000, 2000 and 5000 μg/L). Percent accuracy was evaluated as percent mean of observed concentration from mass spectrophotometer upon spiked concentration injected in the LC-MS/MS. Inter- and Intra-day method precision were obtained from three doxorubicin standard concentrations (5, 50, and 500 μg/L) and presented as percentage of relative standard (% RSD). Intra-day precision was determined by analysing each concentration in duplicate at three time points within a day. Inter-day precision was evaluated by repeating the analysis of all the three mentioned concentrations on three consecutive days.

### Microinjection of zebrafish larvae for generation of breast cancer xenograft

Zebrafish larvae were dechorionated at 2 dpf using fine forceps (Jewelers forceps Dumont No.5, Sigma-Aldrich) under stereo microscope and anesthetized in 0.002% MS-222 (tricaine methanesulfonate) (E10521-10G, Sigma-Aldrich) in 1 X E3 media with 0.2 mM PTU for 1 h in BOD incubator at 28 ⁰C. Anesthetized larvae were mounted laterally in 0.3% agarose on a glass slide. Injection needles were prepared from glass capillaries (TW100-4, World Precision Instruments) pulled using micropipette puller (Sutter Instrument Model P-1000, USA). The inner dimension of the needle tip was maintained 18 μm. 200-400 Vybrant Dil labelled MDA-MB-231 cells were injected into the yolk sac of the mounted larvae. Injected larvae were washed and kept in 1 X E3 media with 0.2 mM PTU at 34 ⁰C in the BOD incubator in dark condition. After 24 h, injected larvae were screened under 1X73 Olympus fluorescence microscope for proper injection and grafting. Larvae with similar sized tumors and no cancer cells in circulation were taken for further study. Success rate of injection which is defined as successful engraftment and enlargement of tumor was calculated as {(Percent larvae showing increased tumor at 96-hour post injection (hpi) compared to 24 hpi in control X Total no. of xenografts in control and treated group)/Total no. of larvae injected} X 100.

### Imaging of zebrafish xenografts

For imaging, zebrafish xenografts were anaesthetised in 0.002% tricaine in 1 X E3 media and mounted in 0.3% agarose on a glass slide. Z-stack fluorescent images in the RFP channel were taken for tumor analysis of the yolk sac region at 10 X magnification using Evos FL auto fluorescence microscope (Life Technologies, Thermofischer scientific). Images were taken at 24- and 96-hpi.

### Qualitative analysis of tumor growth

Doxorubicin has an intrinsic fluorescence (Ex/Em: 470/560 nm) which falls in the range of Vybrant Dil dye (Ex/Em: 549/565 nm). Though the fluorescence coming from both the compound is distinguishable with the naked eye, overlapping doxorubicin fluorescence creates a hindrance while calculating tumor volume using Image J software. Due to this, a qualitative approach was taken to access the tumor growth for this study. Tumor growth at 96 hpi was compared with tumor growth at 24 hpi for each xenograft by five different observers. To keep the data unbiased, these observers were selected from biology background with/without experience in zebrafish handling and fluorescence image analysis. Each tumor was categorized as increase, decrease or similar. Percent larvae with increase/decrease/similar tumor was calculated for both control and doxorubicin treated groups separately. All data was plotted using GraphPad prism software version 8.0.2 as mean ± SEM.

### Statistical analysis

All the statistics were performed using the GraphPad Prism software version 8.0.2. All data are presented as mean ± SEM. * denotes P ≤ 0.05, ** denotes P ≤ 0.001 and *** denotes P ≤ 0.0001.

## Results

### Doxorubicin induced toxicity in zebrafish larvae

For zebrafish xenograft model, it is important that the chemotherapeutic drug is not toxic to the zebrafish itself. Therefore, in order to determine the compatible concentration of doxorubicin in zebrafish larvae we determined the toxicity of doxorubicin in zebrafish larvae. Three days treatment (3-6 dpf) of zebrafish larvae with doxorubicin induced mortality as well as abnormalities at higher concentrations whereas no mortality was observed at lower doxorubicin concentrations (Fig 1). Doses higher than 10 mg/L were toxic to the larvae. The LD_50_ value at 3 dpt was 35.95 mg/L. Percent morphological abnormalities were 51.43% at 50 mg/L doxorubicin concentration whereas no significant abnormalities could be observed in 10 mg/L doxorubicin treated zebrafish larvae. With reference to the mortality data, two doses were taken for doxorubicin uptake studies by LC-MS/MS which includes sublethal doses of 5 mg/L and 10 mg/L. For the doxorubicin treatment of breast cancer zebrafish xenograft, a non-toxic dose of 5 mg/L was used to study its effect in tumor growth.

**Figure 1:**
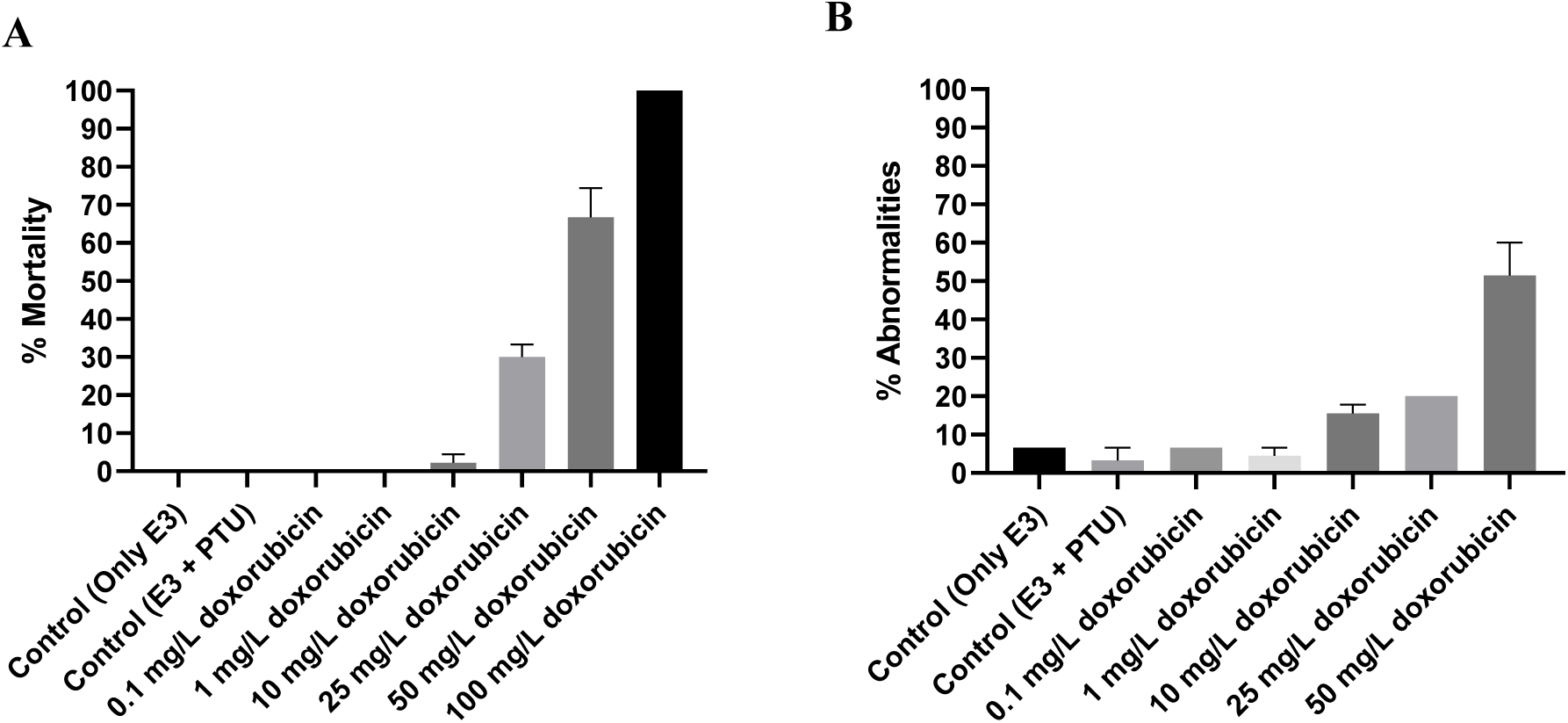
Doxorubicin induces (A) Mortality and (B) Morphological abnormalities at 6 dpf (3 dpt) larvae. Percent mortality was recorded at 3 days of exposure to different concentrations of doxorubicin. Data is plotted from three individual experiments with 5 larvae per treatment group, each performed in triplicates. All data is plotted as mean ± SEM for three experiments containing a total n =15-45 larvae in each group at the start of the experiment.

### HPLC-MS chromatography for doxorubicin detection from zebrafish homogenate spiked with doxorubicin and colchicine

An aspect of HPLC-MS method development is to achieve the retention time within the international standard of ± 0.1 min (19) and fragment ions of each compound that agree with the mass fragmentation pathways. Representative SRM chromatograms of doxorubicin and colchicine are shown in figure 2. Retention time did not show any significant difference for both doxorubicin and colchicine in multiple runs indicating reliable detection of both doxorubicin and colchicine. Fragment ions of doxorubicin were 544.17→361.12, 397.10 m/z which agreed with the mass fragmentation pathways (Fig. 2A) (20,21). Fragment ions of colchicine as internal standard were 400.08→310.16, 358.16 m/z which also satisfies the previous literature (Fig. 2B) (22).

**Figure 2:**
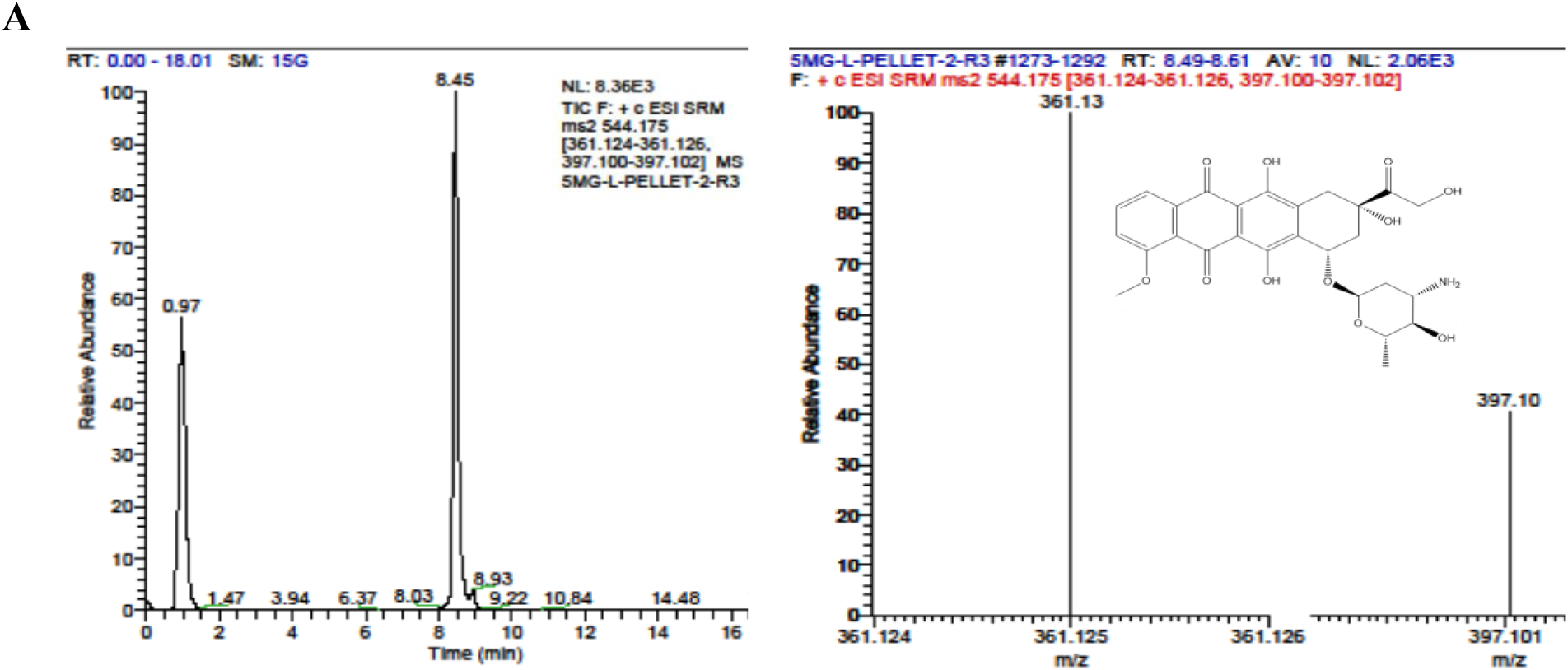
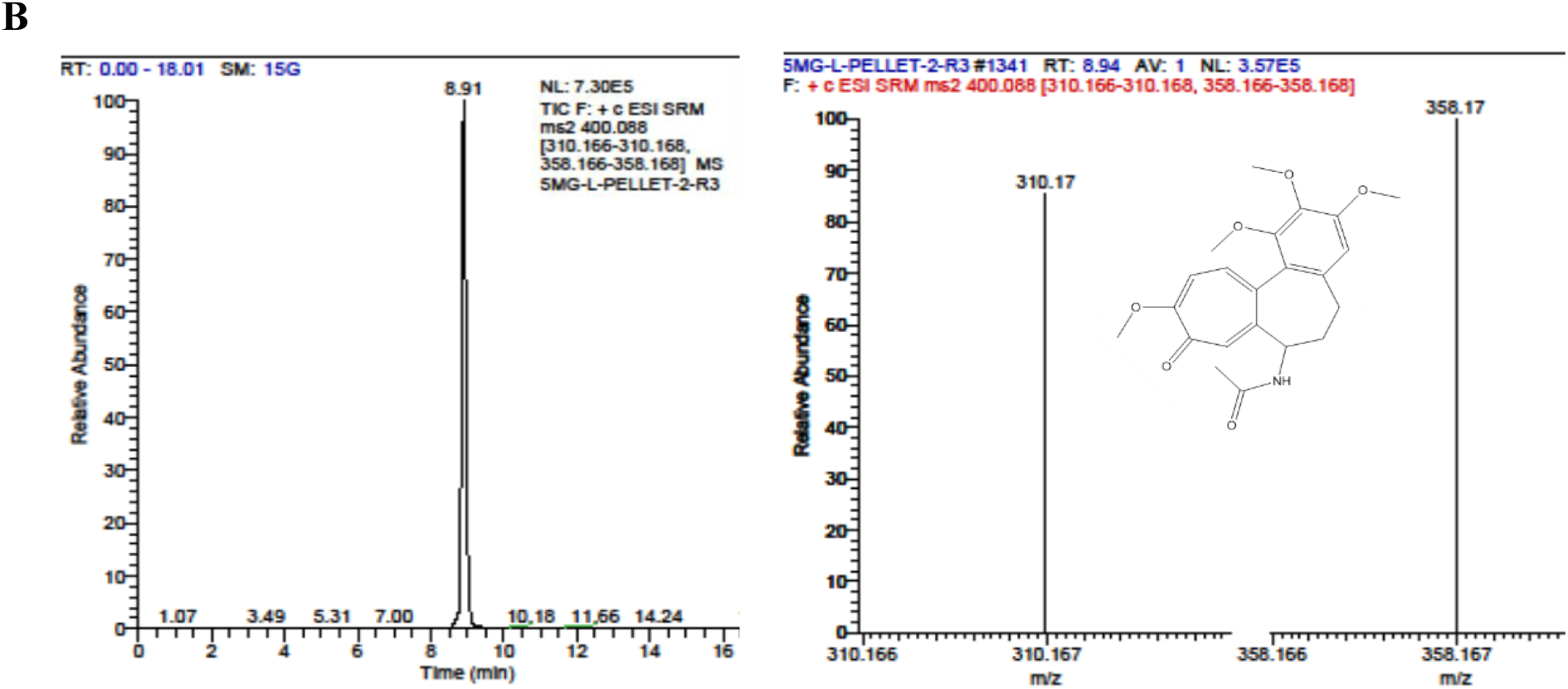
Representative SRM chromatograms for (A) doxorubicin and (B) colchicine. Graphs on the left show retention time and on the right fragment ions of respective compounds. Retention time of both doxorubicin and colchicine from two runs in different sample did not vary more than 0.1 min.

### LC-MS/MS method validation for doxorubicin quantification in zebrafish larval homogenate and fish water

A linear calibration graph for doxorubicin was obtained for concentration range (1-5000 μg/L) with correlation coefficient (R) greater than 0.99. The calibration equation for the graph was y = 5E-05x - 0.0002. The limit of detection (LOD) and limit of quantification (LOQ) are 2 and 5 μg/L respectively and the signal to noise ratio is greater than 10. Intra- and inter-day precision was less than 7% and accuracy was within ± 15% (Table 4).

**Table 4:** Percent accuracy, inter- and intra-day precision of doxorubicin in zebrafish larvae.

| Parameter | Drug dose ($\mu\text{g/L}$) | Doxorubicin |
| --- | --- | --- |
| Accuracy (%) | 5 | 107.85 |
|  | 50 | 92.545 |
|  | 500 | 92.1 |
| Intra-day Precision (% RSD) | 5 | 2.13 |
|  | 50 | 2.25 |
|  | 500 | 1.21 |
| Inter-day Precision (% RSD) | 5 | 2.33 |
|  | 50 | 6.9 |
|  | 500 | 2.18 |

### Determination of doxorubicin uptake in zebrafish larvae

Doxorubicin was quantified in both treatment media (supernatant) and larval pellet. While negligible values were noted for both control samples (supernatant and pellet), the average observed concentration in supernatant of 5,000 and 10,000 μg/L doxorubicin incubated samples were 2709.213 and 5743.7985 μg/L respectively. Calculated doxorubicin concentration per larva in 5,000 and 10,000 μg/L treatment groups was 1.05 and 2.64 ng (Table 5). Pellet 2 was considered as instrumentation error and excluded from analysis.

**Table 5:** Doxorubicin quantification in zebrafish larvae and treatment media.

| Sample | Doxorubicin ($\mu\text{g/L}$) | | Doxorubicin amount/larva (ng) |
| --- | --- | --- | --- |
|  | Initial incubation | Mass spec value |  |
| Supernatant-1 | 0 | 1.01 | NA |
| **Supernatant-2** | 5000 | 2687.327 | NA |
| **Supernatant-3** | 5000 | 2731.099 | NA |
| **Supernatant-4** | 10000 | 5560.576 | NA |
| **Supernatant-5** | 10000 | 5927.021 | NA |
| **Pellet-1** | 0 | 1.147 | 0.019 |
| **Pellet-2** | 5000 | 6176.189 | 102.9364 |
| **Pellet-3** | 5000 | 62.988 | 1.0498 |
| **Pellet-4** | 10000 | 158.206 | 2.6367 |
| **Pellet-5** | 10000 | 158.188 | 2.6364 |

### Doxorubicin treatment reduced tumor growth in MDA-MB-231 breast cancer zebrafish xenografts

Breast cancer zebrafish xenograft was developed with an overall success rate of 20.8% pooled from four individual injections (Fig 3). Inhibitory drug response was observed against doxorubicin from all experimental replicates. Qualitative analysis of tumor growth showed that 43.03% of the control larvae had more tumor increase compared to the treated larvae at 3 dpt. Where, 47.81% of doxorubicin treated larvae had more tumor reduction compared to the control. About 13.43% and 7.78% of control and treated larvae respectively showed no change in tumor size (Fig 4B).

**Figure 3:**
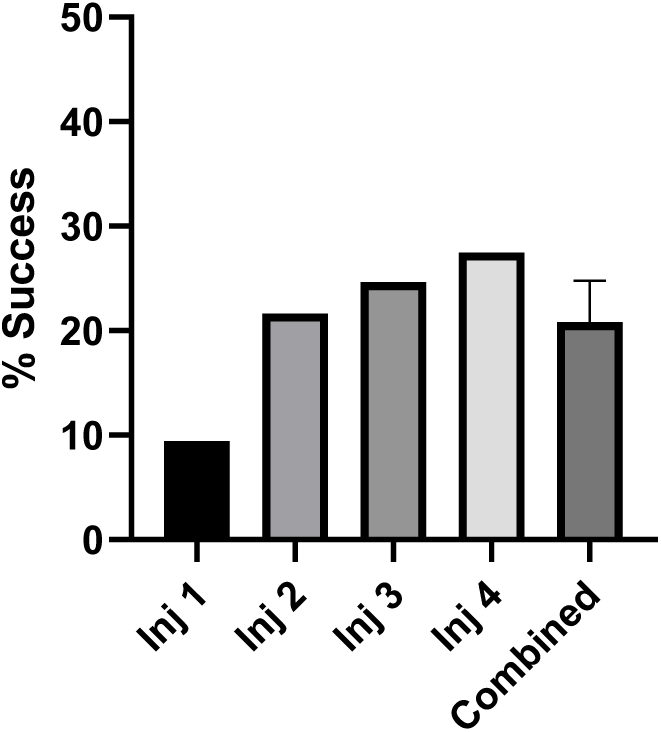
Graph representing success rate of xenograft development by microinjection. Success rate of four different injections (Inj-1, 2, 3, 4) is given along with an overall success rate. Combined success rate is presented as mean ± SEM for all four injections.

**Figure 4:**
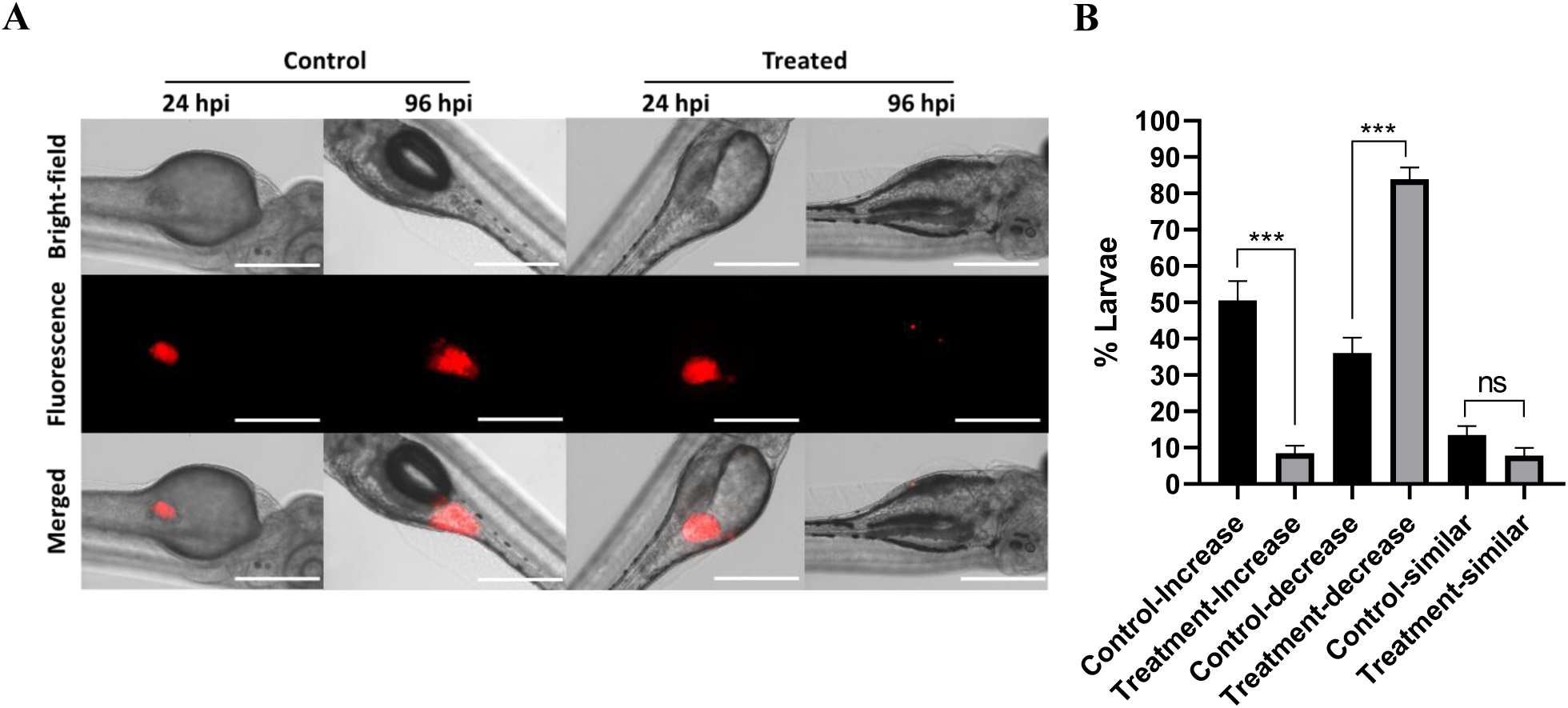
Doxorubicin reduced tumor in breast cancer zebrafish xenografts. (A) Representative images of control and drug treated xenografts. Images are taken at 24- and 96- hpi at 10 X magnification. Treatment of doxorubicin (5 mg/L) is given after 24 hpi imaging. Scale denotes 400 μm. (B) Graph represents qualitative analysis of the tumor growth. All data is plotted as mean ± SEM for four experiments containing a total n = 40 larvae in both control and treatment groups. Unpaired t test was used to analyse the data. *** denotes P ≤ 0.0001.

## Discussion

Zebrafish larvae (3 dpf) when treated with different doses of doxorubicin for 3 days exhibited a dose dependent mortality with an LD_50_ value of 35.95 mg/L at the end of treatment (Fig 1A). Our toxicity data showed mortality starting from 10 mg/L and reached up to 66.67% at 50 mg/L doxorubicin concentration. This data is similar with an earlier study by Chang et al., 2014 where 4 days doxorubicin treatment of 3 dpf larvae had an LD_50_ value of about 35 mg/L and exhibited about 73% mortality at 50 mg/L doxorubicin (23). Based on our toxicity assessment, a non-lethal drug concentration of 5 mg/L was selected for anti-cancer treatment in breast cancer zebrafish xenograft larvae.

LC-MS/MS method validation was completed by following the guidelines for validation of bioanalytical method which requires correlation coefficient of calibration curves to be ≥ 0.99 and intra-and inter-day precision value of ± 15% (24,25) as mentioned in table 4. HPLC-MS/MS with SRM mode used in our method provides accurate analysis for quantification. Quantification of doxorubicin in the zebrafish larval homogenate revealed body burden of 1.05 and 2.64 ng/larvae in 5,000 and 10,000 μg/L treatment groups respectively (Table 5). Here, we observed an increase in the drug uptake per larva with increasing treatment concentration a trend usually observed for other compounds as well (17,18). Additionally, doxorubicin in the treatment media was detected at 2709.213 and 5743.7985 μg/L respectively (Table 5). Here, the observed concentration was lesser than the added concentration which could be because of degradation of doxorubicin over the incubation period. Moreover, doxorubicin has been reported to degrade in physiological pH (26) and as well as with increasing temperature (27). Reduced concentration is also detected for other compounds when subjected to LC-MS/MS (28).

Successful engraftment of melanoma cells in blastula stage of zebrafish was first reported in 2005 (29), after which the feasibility of the procedure was performed by multiple researchers (30,31). Different parameters such as percent survival (32) and percent engraftment (33) of zebrafish xenografts were accessed by different groups. While percent engraftment for three different pancreatic cells in zebrafish larvae was given by Wang et al.,2022, they didn’t provide the method for calculation (33). Hence, there remained a gap in the assessment of injection efficiency. This is the first study in zebrafish xenograft defining success rate where we provide a definitive method of success rate calculation which will determine the efficiency of injection. Success rate was determined to be 20.8% in our experiments when calculated for our MDA-MB-231 breast cancer zebrafish xenograft. Although similar approach for success rate calculation was not employed previously, we believe this value is good based on the previous literature where only few injected larvae were seen to have tumor growth even after screening the larvae for experiments (34,35). Further, a significant decrease in tumor growth was observed through qualitative analysis, when the zebrafish xenografts were treated with doxorubicin (Fig. 4B). Approximately 47.81% (Fig. 4B.bar 3 and 4) of the treated group had more tumor reduction compared to control at 3 dpt which confirms doxorubicin’s anti-cancerous effect. Furthermore, our qualitative findings align with the results from previous studies showing tumour reduction in MDA-MB-231 zebrafish xenografts treated with 4.3 mg/L doxorubicin (10). From the LC-MS/MS quantification results we can further infer that the tumor reduction in larvae is caused by 1.05 ng of doxorubicin which is the body burden per larvae when treated with 5,000 μg/L doxorubicin. This reference value of doxorubicin can be used by researchers to validate their MDA-MB-231 zebrafish xenograft model in future perhaps by directly injecting doxorubicin.

In summary, our study establishes an important methodology for assessing doxorubicin uptake in zebrafish larvae and provides a reference quantity for its anti-cancerous effect. Validation of our LC-MS/MS quantification method through the provided guidelines ensured precise analysis.

Further, our developed method can be perhaps used to accurately quantify doxorubicin amount needed to develop other disease models such as the widely used doxorubicin induced zebrafish cardiomyopathy model (36). It may also offer a high throughput platform to measure and validate the increased uptake of doxorubicin formulations designed to minimize its side effects (16,37).

## Acknowledgment

This work was supported by the IIT Hyderabad, ICMR-DHR Adhoc and Japan friendship 2.0 grants to AB. Ministry of Education fellowship for PhD to GR, UGC-JRF fellowship for PhD to AP. The authors thank Centre for Mass Spectrometry, CSIR-Indian Institute of Chemical Technology, Hyderabad for providing their Mass Spectrometry facility.

## Statement and declaration

### Ethics approval

All animal handling and experiments were conducted in accordance with the guidelines prescribed by the Committee for Control and Supervision of Experiments on Animals (CCSEA), Government of India.

### Consent to participate

Not applicable

### Consent to publish

Not applicable

### Availability of data and material

All the data are available upon request to the corresponding author

### Credit authorship contribution statement

**GR:** Experiments, Analysis, Drafting, Editing; **AP**: Experiments; **SD:** Imaging; **AR:** Imaging;

**AB:** Conceptualization, Analysis, Drafting, Editing, Reviewing, Funding acquisition.

